# Mechanical flexibility reduces the foreign body response to long-term implanted microelectrodes in rabbit cortex

**DOI:** 10.1101/058982

**Authors:** H S Sohal, G J Clowry, A Jackson, A O’Neill, S N Baker

**Affiliations:** Media Lab, McGovern Institute, Microsystems Technology Laboratories, MIT, Cambridge, MA, 02139, USA; Institute of Neuroscience, Newcastle University, Newcastle Upon Tyne, NE2 4HH, UK; School of Electrical and Electronic Engineering, Newcastle University, Newcastle Upon Tyne, NE1 7RU, UK

**Author notes:** = *these authors contributed equally to this work* (A Jackson, A O’Neill, S N Baker).

## Abstract

Micromotion between the brain and implanted electrodes is a major contributor to the failure of invasive microelectrodes. Movements of the electrode tip cause recording instabilities while spike amplitudes decline over the weeks/months post-implantation due to glial cell activation caused by sustained mechanical trauma. We compared the glial response over a 26-96 week period following implantation in the rabbit cortex of microwires and a novel flexible electrode. Horizontal sections were used to obtain a depth profile of the radial distribution of microglia, astrocytes and neurofilament. We found that the flexible electrode was associated with decreased gliosis compared to the microwires over these long indwelling periods. This was in part due to a decrease in overall microgliosis and enhanced neuronal density around the flexible probe, especially at longer periods of implantation.

## 1. Introduction

Penetrating electrode arrays offer the ability to monitor neural activity in a wide variety of experimental paradigms and in clinical settings, at single-cell resolution. However, when such arrays are implanted chronically, the brain immune response is activated, leading to the degradation of viable signal from neurons and recording instabilities [1,2]. Regardless of the electrode design (whether silicon or microwire electrodes), there is a clear activation of resident microglia and infilitrating macrophages due to blood-brain-barrier damage around the implants ranging up to 100 microns, accompanied by reduced neural density after periods as short as 4 weeks [3–6]. Furthermore, hypertrophic astrocytes surrounding the implant from 100-300 microns can result in a high impedance barrier between the electrode and the surrounding brain tissue [3]. The lack of neurons, high impedance glial scar and degradation of the electrode due to hostile immune factor release contribute to generalize electrode failure.

Over the past 60 years, many factors have been identified that can cause an enhanced foreign body response to implanted electrodes: Damage of vasculature upon initial insertion leading to decreased or no blood perfusion [7] for the brain tissue where the electrode resides [8–10], damage to the blood brain barrier causing infiltration of peripheral immune factors [7,11] and activation of microglia and astrocytes releasing hostile factors which compromise neuronal function [3,12] and electrode materials [13,14]. This persisting immune response is likely caused by micromotion induced trauma [15]. Typical electrode materials include silicon and microwires, which have a higher Young’s modulus (150 GPa) than brain tissue (5-30 KPa), meaning that microelectrodes anchored to the skull cannot accommodate movement of the brain within the cranial cavity. Even if electrodes are floating with the brain, the deformation of the tissue will change the position of the deep neurons relative to the anchoring point at the brain. This inability to deform with the brain persistently irritates the surrounding brain tissue and can cause a heightened detrimental immune response.

One way to mitigate the modulus mismatch between the brain and electrode materials, is to use materials with a lower Young’s modulus, for example Parylene-C or Polyimide insulated thin film electrodes that have been shown to record for periods up to 4-12 weeks [16–18]. However, to find a clinically viable solution for the instabilities in Brain machine interfaces it is important to consider the foreign body response over longer indwelling periods.

Here we present a comparison of the foreign body response (FBR) to novel flexible electrodes (‘Sinusoidal probe’) and to conventional microwire electrodes over very long indwelling periods from 24-96 weeks in the rabbit cortex. We used horizontal sections to obtain a depth profile over each electrode, evaluating microglia, astrocytes and neurofilament to give a three-dimensional assessment of the FBR. Post-mortem histology suggests that overall the flexible electrodes had a reduced response compared to the microwires, with enhanced neural survival. These results are applicable to many flexible designs and validate the need to reduce micromotion-induced trauma to enhance electrode longevity and minimize the immune response.

## 2. Material and methods

### 2.1 Microelectrodes

Details of the flexible sinusoidal probe have been published previously [19]. Briefly, the overall electrode body was 20 μm deep and 35 wide, and 3 or 5.5mm long. The electrode shaft consisted of 10 sinusoidal cycles of 100 amplitude and 500 μm period, and was linked via a 3 cm long and 3 mm wide ribbon cable to a standard connector (micro ps1/ps2 series, Omnetics Connector Corporation, USA). The probe was made out of flexible Parylene-C and had tungsten-titanium conductive metal for the recording sites (96 μm^2^) and conductive tracts. Post-processing, a spheroid polyimide anchor was added to the recording end (100 μm diameter), to anchor the three protruding recording sites. The ‘sinusoidal’ shaft would counteract the motion of the brain, with the recording sites restricted in movement, relative to the recording tissue of interest.

The flexible probe was temporarily attached to a sharp and rigid carrier for insertion (0.229 mm diameter steel electrodes, 2-3 μm tips; Microprobe INC, USA) using poly-ethylene-glycol (PEG, MW. 6000, Sigma Aldrich, USA).

Microwire electrodes (50 μm diameter Teflon-insulated tungsten, Advent, UK) were constructed by appropriately deinsulating one end of the wire and crimping a connector to form reliable electrical contact (Amphenol, USA). The other end was cut-flush, which would be implanted into the rabbit brain.

### 2.2 Animals and Surgery

All experiments were approved by the local ethics committee at Newcastle University and were performed under appropriate Home Office licences in accordance with the UK Animals (Scientific Procedures) Act 1986.

Electrodes (4-5 Sinusoidal probes and 4 Microwire probes) were implanted into the sensorimotor representation of four New Zealand white rabbits *(Oryctolagus cuniculus;* Two Subjects were used for the 26 week characterization and single subjects were used for 52 and 96 week characterization.

After a midline skin incision, electrodes were inserted stereotaxically relative to bregma, between 4mm anterior to 4 mm posterior and 0.5 to 7 mm lateral (Gould, 1986; Swadlow, 1989; 1990; 1994; Swadlow and Hicks, 1996), after burr-hole craniotomies were made in the specific locations. Electrodes were inserted quickly [20] using a stereotaxic manipulator (Kopf, USA). The PEG was dissolved with warm saline to release the electrode from the carrier, which was subsequently removed. Microwire electrodes were inserted manually with hooked surgical forceps. Connectors were attached with skull screws and dental cement. A wire wrapped around one skull screw served as ground and reference for the recordings.Animals were anaesthetized with hypnorm (0.3ml/kg i.m.) and midazolam (2 mg/kg i.v.) prior to transcardial perfusion with phosphate buffered saline (PBS) and then formal saline. The relevant brain regions were removed and transferred to a 10 % sucrose solution. The day before microtome sectioning, the extracted brain was transferred to a 30% wt sucrose solution for cryoprotection. 50 μm thick sections were obtained over the entire electrode profile.

### 2.4 Immunostaining

Horizontal slices corresponding to the top, middle and bottom of the electrode profile were stained for microglia, astrocytes and neurofilament. Slices were incubated with 3% normal horse serum (Vector Labs, UK: S-2000) for 1 hour to prevent non-specific binding. Sectioned slices were incubated overnight with relevant concentrations of the primary antibodies GFAP (Sigma-Aldrich, UK), isolectin-b4 (Vector Labs, UK: Biotinylated Griffonia (Bandeiraea) Simplicifolia Lectin 1, B-1105) and where appropriate, SMI-32 (Cambridge Biosciences, UK: R-500) made up with PBS-triton on a rocker at 4°C (table 1). GFAP and Neurofilament slices underwent incubation with biotinylated antimouse for two hours. All slices were then incubated with horse radish peroxide streptavidin (HRP strep, Vector Labs, UK: SA-5004) for one hour. After every incubation stage three, five minutes washes with PBS were performed on the slices on a rocker.

Finally, the Diaminobenzidine (DAB) reaction was performed. One tablet per 5 ml of PBS of peroxide and urea hydrogen was formulated (Sigma-Aldrich, UK). Slices were incubated with DAB for 5 minutes before being transferred into PBS filled wells. Slices were mounted on gelatin-coated slides according to electrode profile and staining and were left to dry overnight. The slices then underwent a series of alcohol (5 minutes of 70%, 95%, 100%, 100%) and two, 10 minute histoclear washes (Sigma-Aldrich, UK), before cover slips were mounted with histomount (Sigma-Aldrich, UK).

**Table 1.**
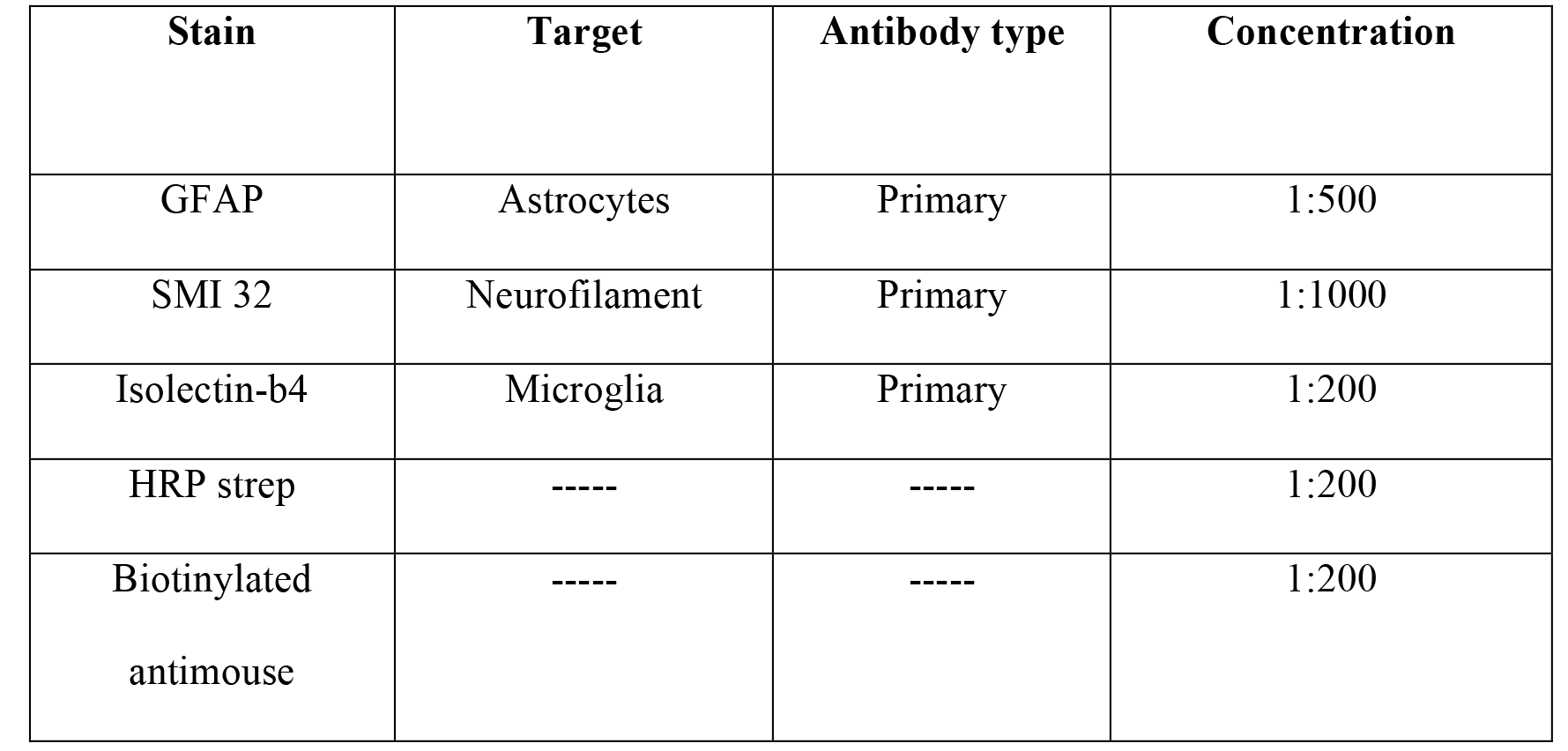
Antibodies and relative concentrations used for analysing the immune response around electrode implants. All antibodies were diluted in PBS-0.3% triton.

| Stain | Target | Antibody type | Concentration |
| --- | --- | --- | --- |
| GFAP | Astrocytes | Primary | 1:500 |
| SMI 32 | Neurofilament | Primary | 1:1000 |
| Isolectin-b4 | Microglia | Primary | 1:200 |
| HRP strep | ----- | ----- | 1:200 |
| Biotinylated<br>antimouse | ----- | ----- | 1:200 |

### 2.5 Analysis

All statistical comparisons were between multiple electrodes within the same animal to avoid inter-individual variability. For immunohistochemisty, slices were examined under an optical microscope at x10 magnification and images taken of electrode tracks and background at a pre-saturation exposure time dependent on the staining with the use of axiovision software (Carl Zeiss Microimaging, Germany). Each image was then normalised in accordance to endogenous background staining in the Matlab environment (2009a, MathWorks, USA) and inverted. Background images were taken at least 1 mm away from the electrode implantation site. Overall background intensity values were subtracted from electrode tract images to leave the glial response around electrodes. Images were imported into ImageJ64 software (NIH, USA) and the radial distribution intensity taken for slices from microwire and sinusoidal probes, centred on the implant sites and according to the electrode shaft profile.

This analysis was carried out on all sections for microglia, astrocytes and where appropriate neurofilament. Normalised integrated intensity and distance from the electrode values were obtained for each slice for all identifiable electrode tracks at the three electrode shaft profiles (top, middle and bottom). To analyse the overall effect of the electrode implants, the mean of at least 15 electrode implantation sites for each depth profile were compared between the two electrode types. An overall effect was analysed as gliosis may change from one electrode implant site to another (Rousche and Normann, 1998).

Further analysis was then carried out in the Matlab environment (2009a, MathWorks, USA), where paired t-tests were performed comparing both electrode types on specific intensity values corresponding to 50-500 away from the electrode implantation centre across all electrode tracts and profile depths, at 50 spacing. This was in accordance to literature to allow for comparison purposes (Winslow et al., 2010a, Winslow and Tresco, 2010, Potter et al., 2012). Distance comparisons were also corrected to account for potential differing implantation site sizes, on a per-electrode tract basis.

To account for multiple comparison error, the Bonferroni correction was performed. The smallest p-value obtained for each depth profile comparison had to be smaller than 0.05 divided by the total number of t-tests performed (10) to account for multiple comparison error. If this was the case, all t-tests performed were accepted for the specific depth profile comparison.

The probability of a false positive (type I error) being reported without the Bonferroni correction is given by 1-(1-α)^n^, where α is the confidence interval and n is the number of t-tests performed. For our study this gives a value of 0.40.

This analysis process is summarized in Fig. 1.

**Figure 1.**
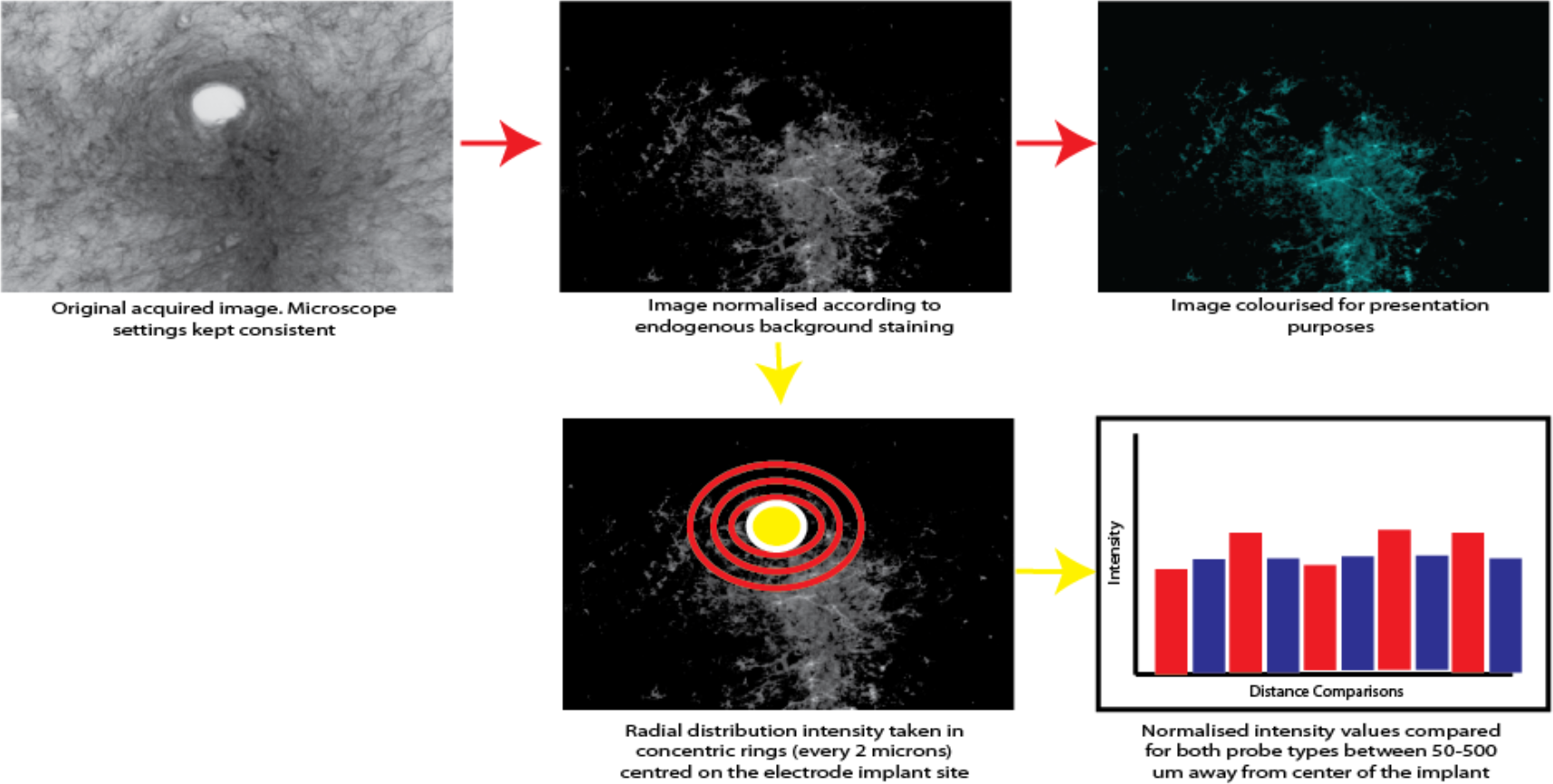
The sequential steps for glial response normilisation and subsequent statistical analysis using a radial distribution to compare staining intensity for microgliosis, astrcytosis and neuralfilament staining.

## 3 Results

### 3.1 Top profile comparison

There was a significant reduction in the astrocytic response (Fig. 2) observed up to 300 μm and 250 μm, away from the electrode implant for the sinusoidal probe at 52 (Fig 2A,B) and 96 (Fig. 2E,F) weeks, respectively.

**Figure 2.**
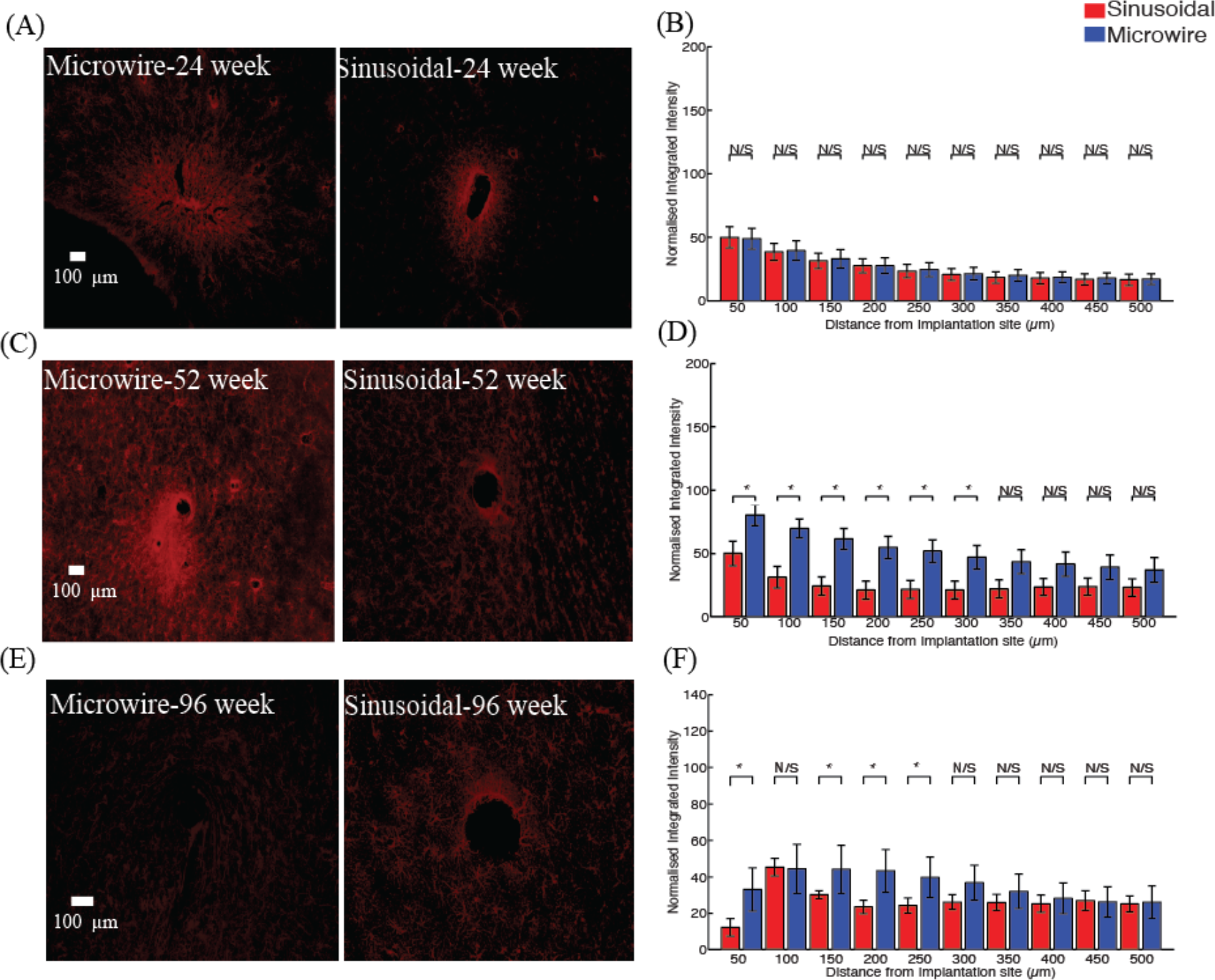
The astrocytic response for the top profile for the sinusoidal probe and microwire probe for 26 (A,B), 52 (C,D) and 96 (E,F) week time points. Reduction in astrcytosis was found at 52 and 96 weeks for the flexible probe

The microglial reaction (Fig. 3) was reduced between 200-500 μm and 0-250 μm for 52 (Fig. 3A,B) and 96 (Fig. 3C,D) week time points.

**Figure 3.**
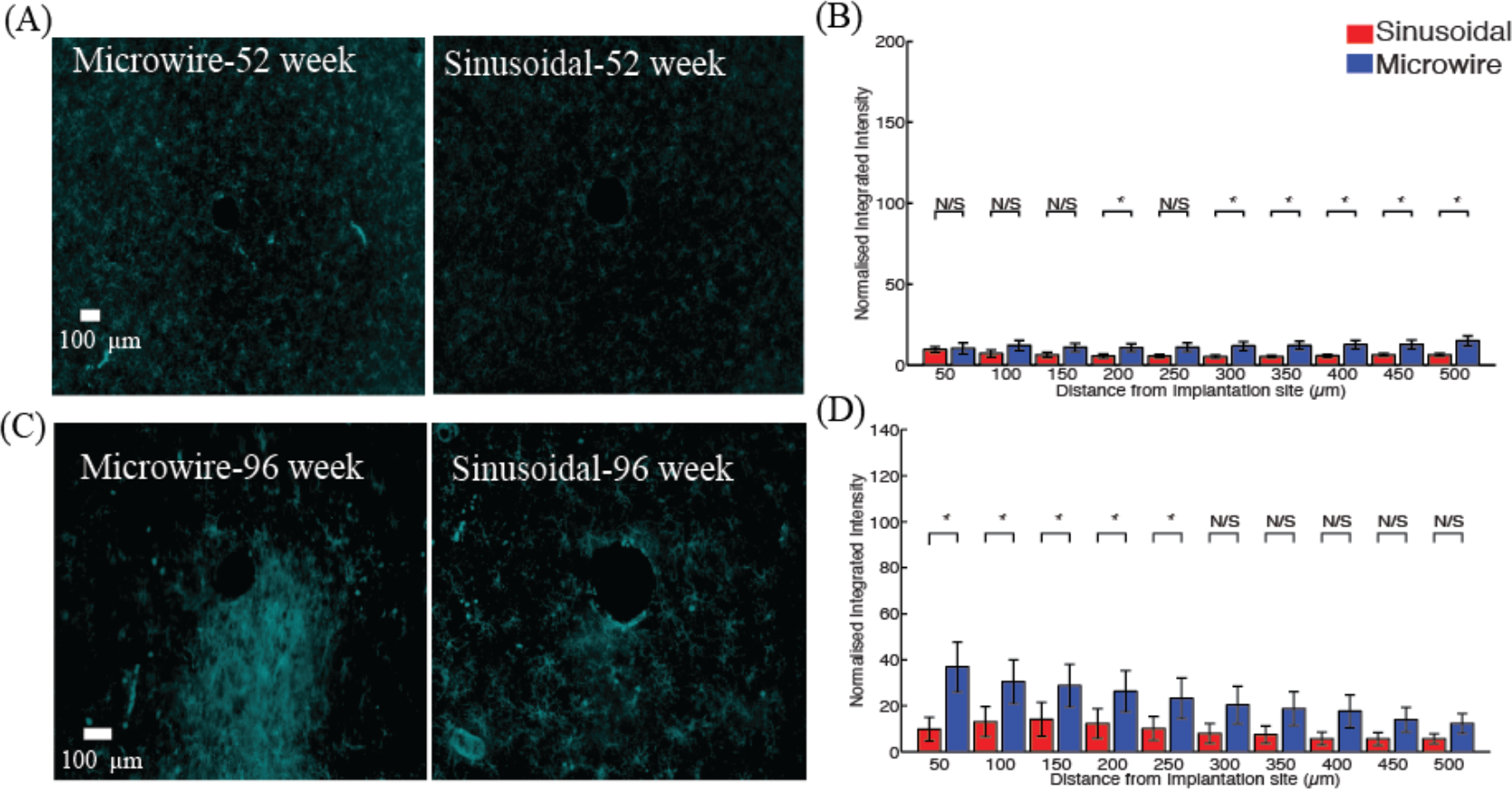
The microglial response for the top profile for the sinusoidal probe and microwire probe for 52 (A,B) and 96 (C,D) week time points. Reduction in microgliosis was found at 52 and 96 weeks for the flexible probe.

An increase in neurofilament (Fig. 4) was observed up to 150 and 50 away for 52 (Fig. 4A,B) and 96 (Fig. 4C,D) week time points. The reduction in the microglial response was often accompanied by an increase in neurofilament staining around the flexible probe.

**Figure 4.**
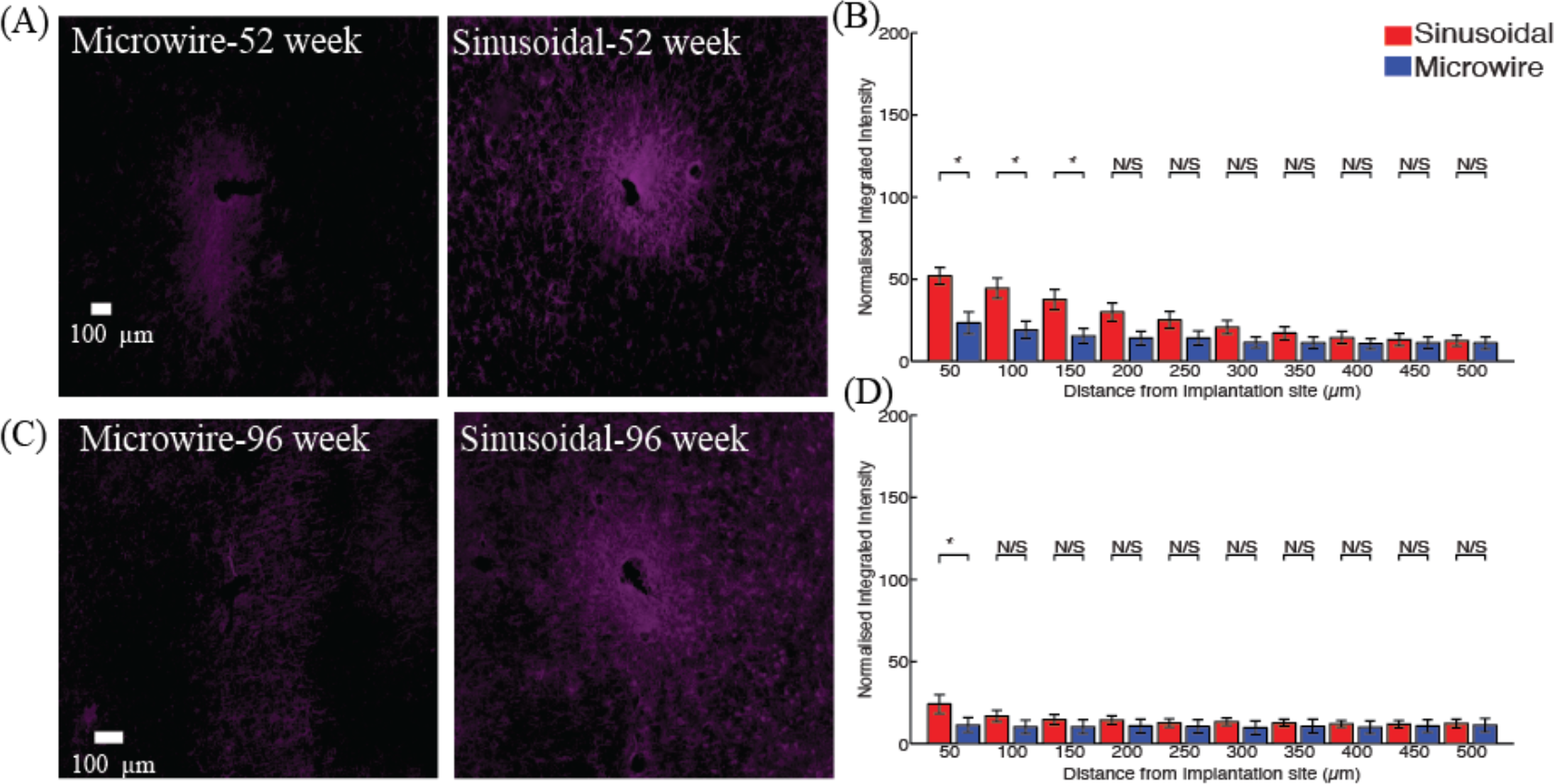
Neuronal density for the top profile for the sinusoidal probe and microwire probe for 52 (A,B) and 96 (C,D) week time points. Increased neurafilament was found at 52 and 96 weeks for the flexible probe close to the probe.Middle profile comparison

### 3.2 Middle profile comparison

There was a significant reduction in the astrocytic response (Fig. 5) observed up to 200 and 100 μm away from the electrode implant for the sinusoidal probe for 52 (Fig. 5A,B) and 96 (Fig. 5E,F) weeks, respectively. The microglial reaction was similar for both electrode types (Fig. 6).

**Figure 5.**
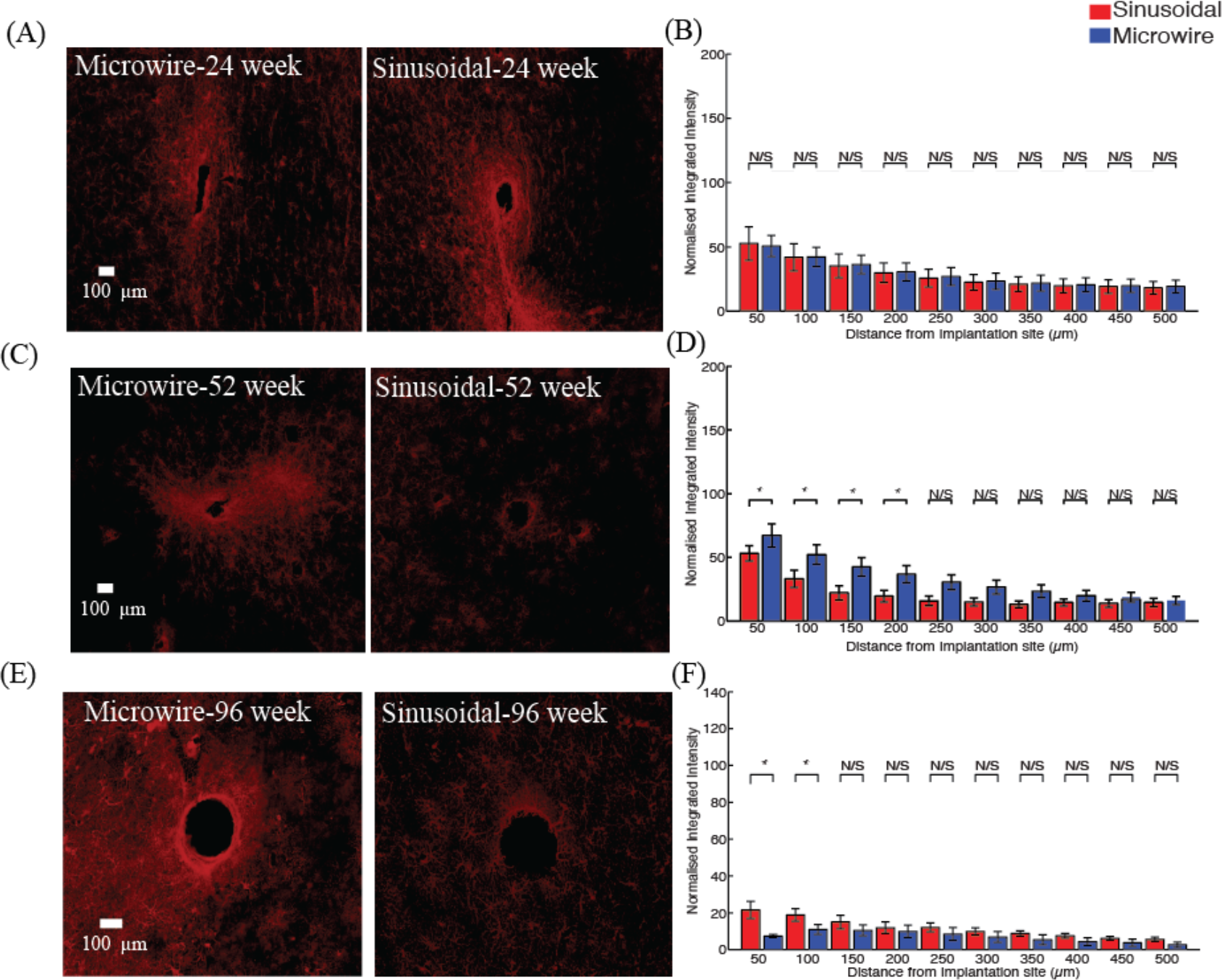
Astrocytic response for the middle profile for the sinusoidal probe and microwire probe for 26 (A,B), 52 (C,D) and 96 (E,F) week time points. Decreased astrocytosis was found at 52 and 96 weeks for the flexible probe close to the probe.

**Figure 6.**
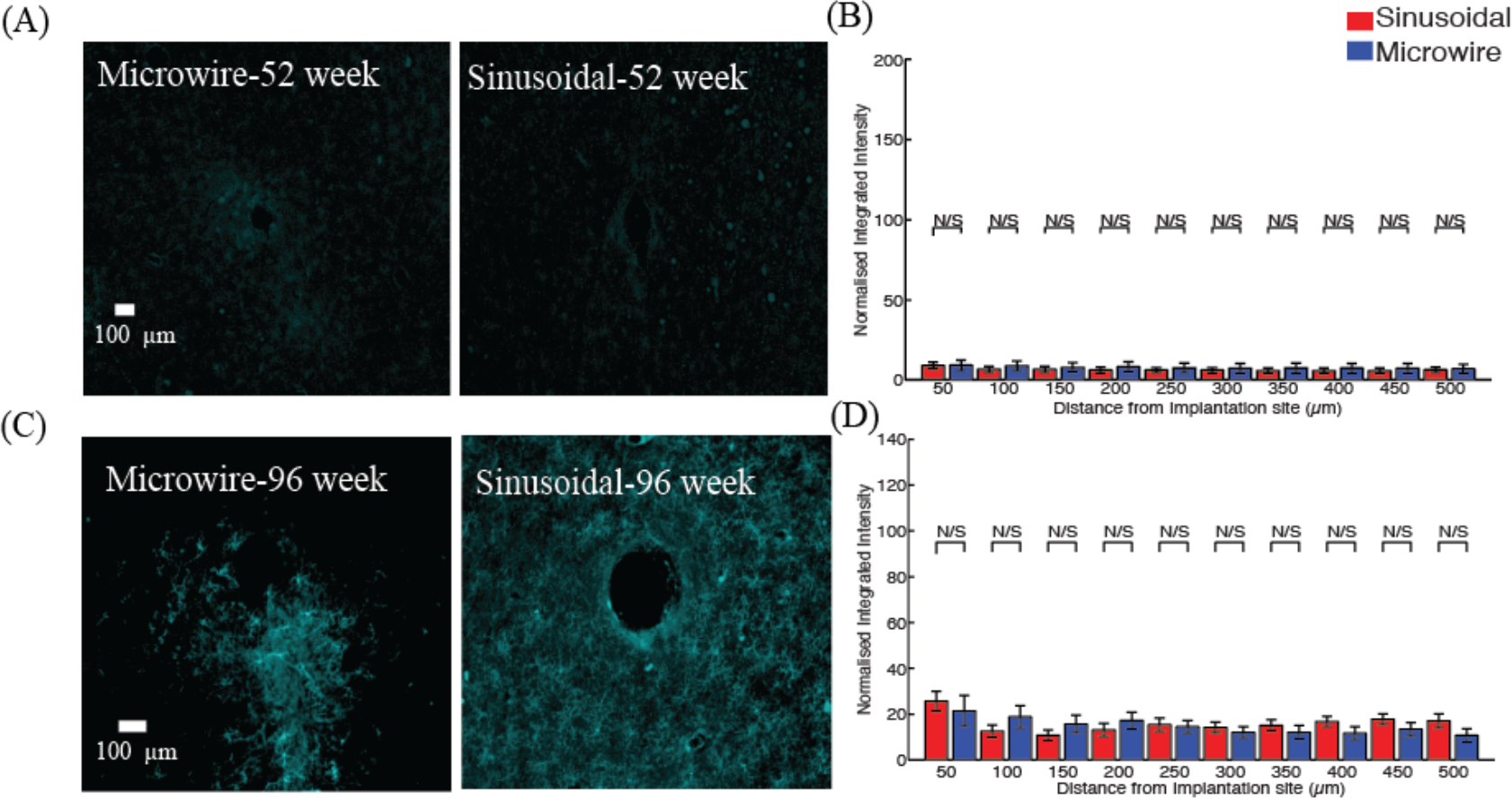
Microglial response for the middle profile for the sinusoidal probe and microwire probe for 52 (A,B) and 96 (C,D) week time points. Similar responses were found for both probe types.

An increase in neurofilament (Fig.7) was observed up to 150 and 100 away for 52 (Fig. 7A,B) and 96 (Fig. 7C,D) week time points.

**Figure 7.**
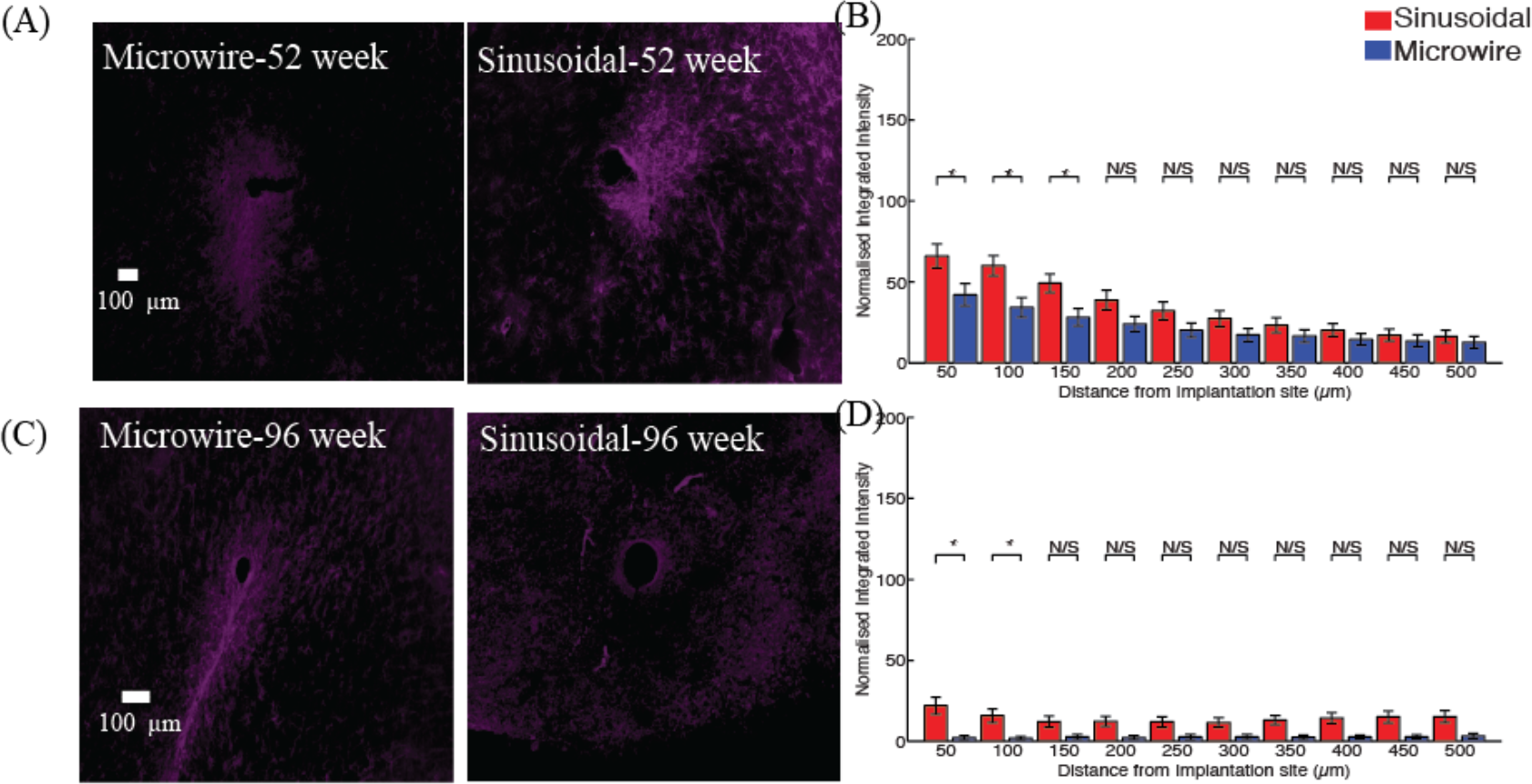
Neuronal density for the middle profile for the sinusoidal probe and microwire probe for 52 (A,B) and 96 (C,D) week time points. Increased neurafilament was found at 52 and 96 weeks for the flexible probe close to the probe.
3.3 Bottom profile comparison

### 3.3 Bottom profile comparison

A reduction in the astrocytic response was observed up to 250 and 450 μm away from the electrode implant for the sinusoidal probe for 24 (Fig. 8A,B) and 96 (Fig. 8E,F) week, respectively.

**Figure 8.**
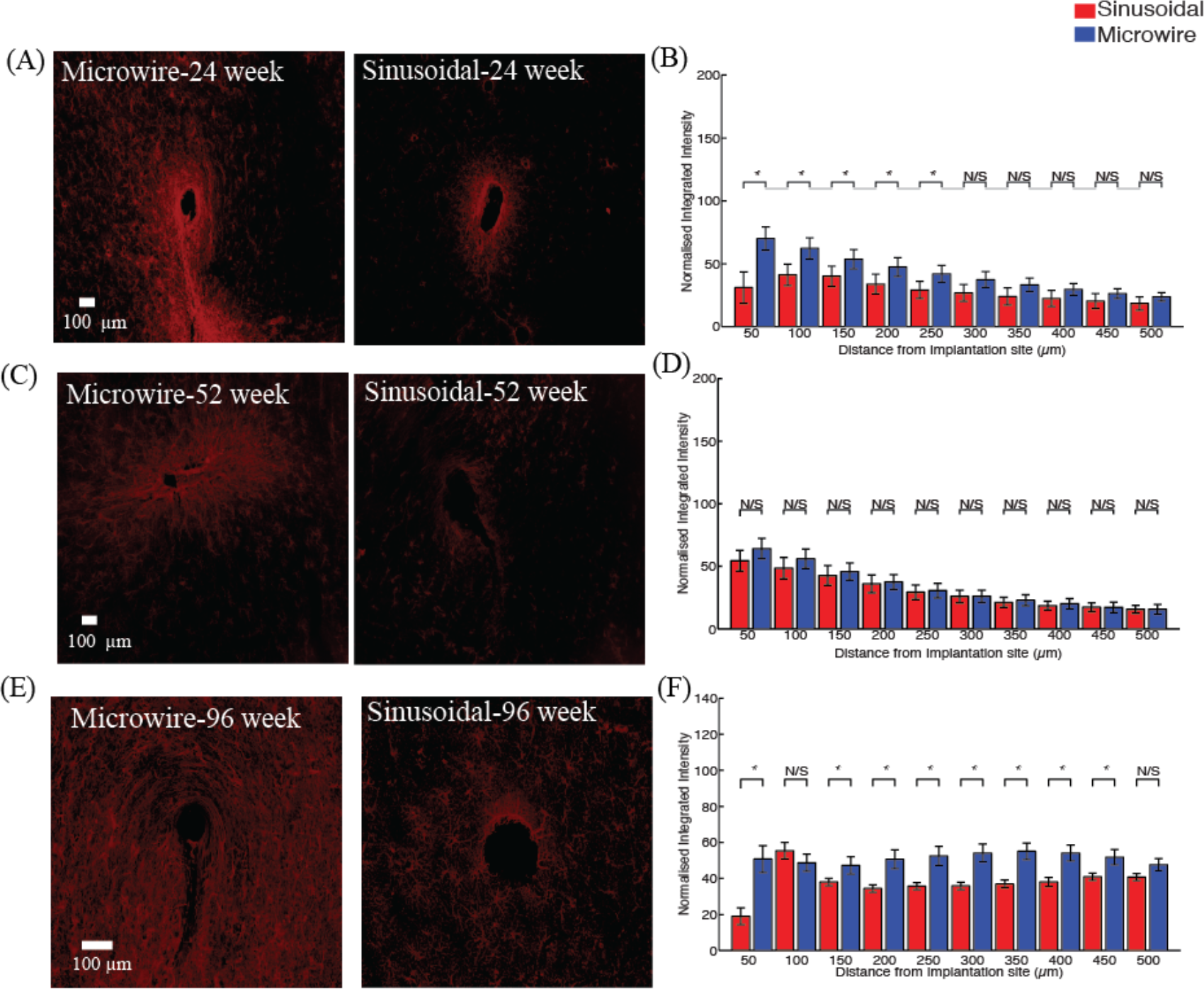
Astrocytic response for the tip region for the sinusoidal probe and microwire probe for 26 (A,B), 52 (C,D) and 96 (E,F) week time points. Decreased astrocytosis was found at 24 and 96 weeks for the flexible probe.

The microglial reaction was reduced up to 500 μm away for both 52 (Fig. 9A,B) and 96 week time points.

**Figure 9.**
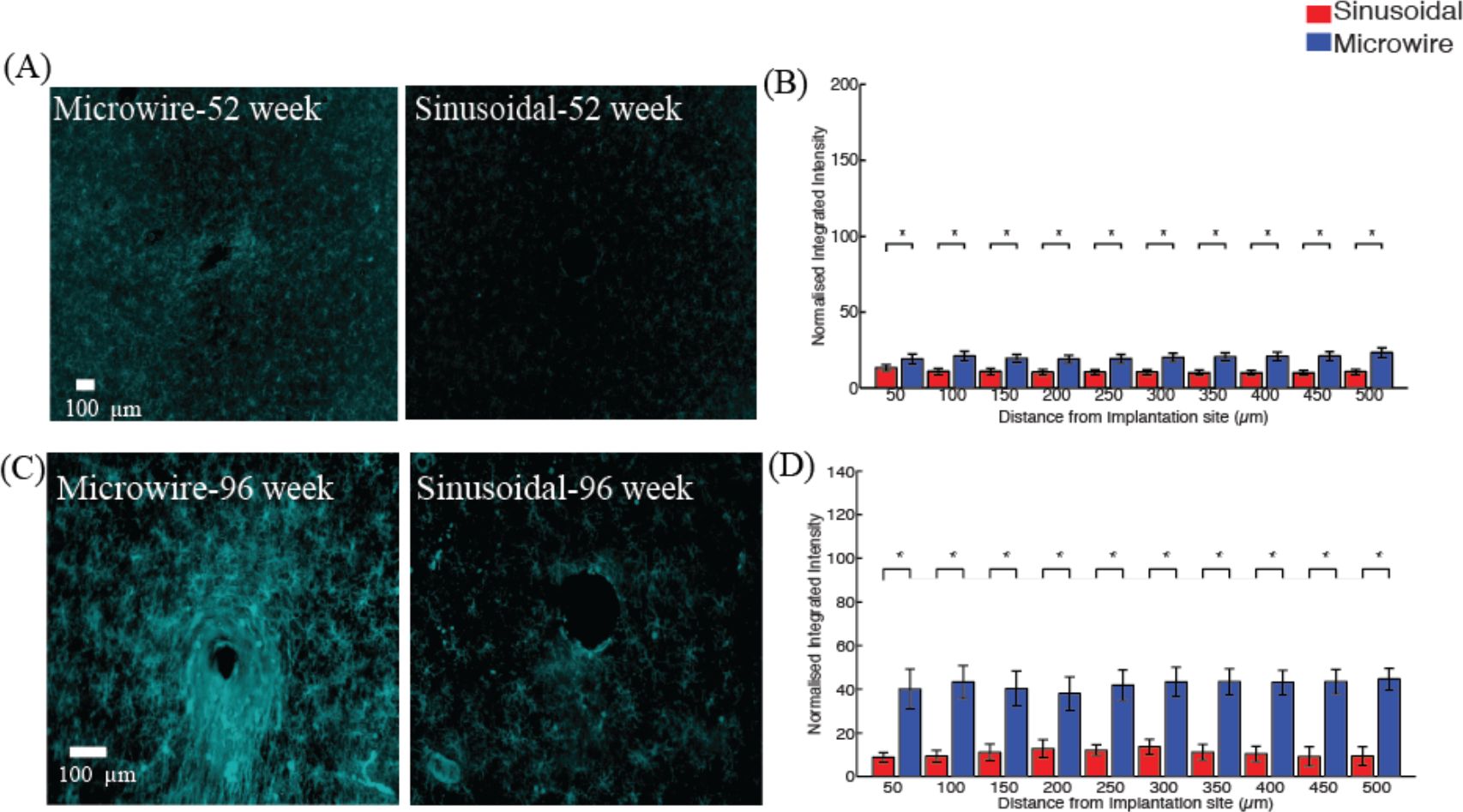
Microglial response for the tip region for the sinusoidal probe and microwire probe for 52 (A,B) and 96 (C,D) week time points. Decreased microgliosis was found at 52 and 96 weeks for the flexible probe at all the distance comparisons.

An increase in neurofilament was observed up to 500 and 100 μm away for 52 (Fig. 10A,B) and 96 (Fig. 10 C,D) week time points. The increase in neurofilament was related to a decrease in the microglial response around the probe’s recording sites, showing the viability of the tissue around the flexible probe more than 52 week post-implant.

**Figure 10.**
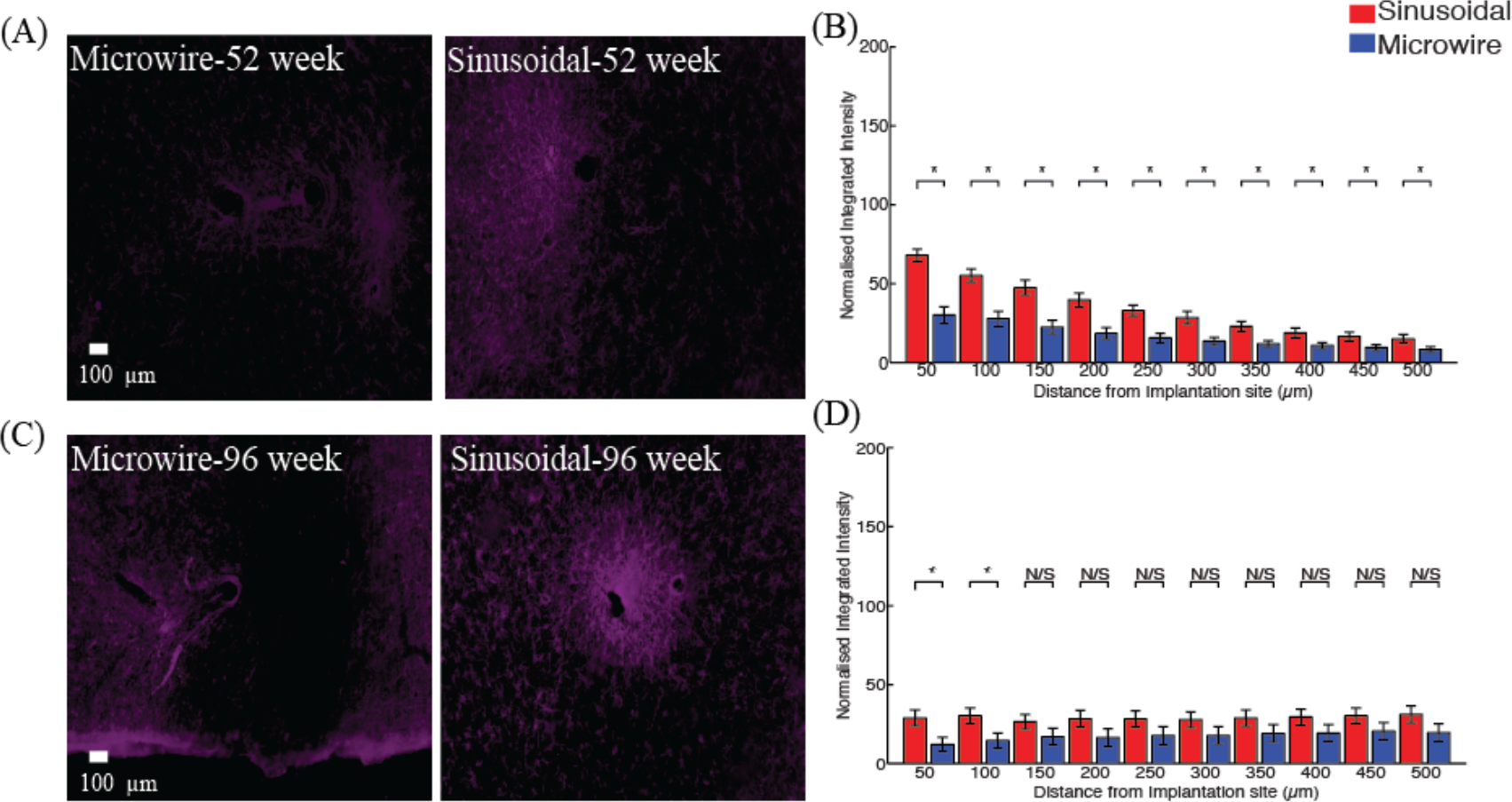
Neuronal density for the tip region for the sinusoidal probe and microwire probe for 52 (A,B) and 96 (C,D) week time points. Increased neuronal density was found at 52 and 96 weeks for the flexible probe.

### 3.4 Summary of results

A summary of the histological comparisons is provided in table 2.

**Table 2.** Summary of histological comparisons at 24,52 and 96 week chronic indwelling points for both microwire and sinusoidal probeGeneral implantation footprint

| Timepoint | Top profile | Middle profile | Bottom profile |
| --- | --- | --- | --- |
| 26 week<br>Astrocytes | _____ | _____ | ↓ |
| 52 week<br>Astrocytes | ↓ | ↓ | _____ |
| 52 week<br>Microglia | ↓ | _____ | ↓ |
| 52 week<br>Neurofilament | ↑ | ↑ | ↑ |
| 96 week<br>Astrocytes | _____ | _____ | ↓ |
| 96 week<br>Microglia | ↓ | _____ | ↓ |
| 96 week<br>Neurofilament | ↑ | ↑ | ↑ |

### 3.5 General implantation footprint

Implant footprints (size of the hole resulting from the electrodes) were also measured for all the profile depths. For the top profile the mean implant size (n=15, ±S.E.M.) was 96.6± 1.4 and 108. 4±10.4 for the microwire and sinusoidal probe, respectively. For the middle profile the mean implant size (n=15) was 120.2± 15.0 and 146.1± 19.7μm for the microwire and sinusoidal probe, respectively. For the bottom profile the mean implant size (n=15) was 101.5± 3.1and 147.3± 14.1μm for the microwire and sinusoidal probe, respectively *(t(15)=3.14,P<0.05).* The implant profile for the sinusoidal probe was larger than the microwire for the bottom profile, perhaps owing to the incorporation of the polyimide anchor.

## 6. Discussion

We compared the FBR to a flexible electrode and conventional microwire over very long indwelling times. Overall, there was a decreased FBR for the flexible probe at the differing depths evaluated. Interestingly, decreased microglial reaction was associated with an increase in neural density around the flexible probe at all of the depths measured. For longer time-points, there was an apparent increase in microglia, accompanied by decreased neuronal density around the microwire electrode. This supports existing literature that a decreased microglial reaction should promote neural viability. Interestingly we found a similar microglia response for the middle profile of both electrodes, perhaps owing to the moving shaft that was designed to counteract the brain motion at the electrode tip. However, the response is not worse than the microwire probe.

In this study we used horizontal sections to evaluate the FBR at different depths relating to different parts of the probe. For some tracts, it was noticeable that the sinusoidal probe had a larger implant profile perhaps due to the formation of the polyimide-anchoring ball, which could vary in diameter (100-200 μm). However, even with this discrepancy it is noted that reduced FBR was achieved overall compared to the microwire probe. Further, the implantation used a stiff, sharp carrier for the sinusoidal probe, in contrast to the direct insertion of microwires. Both the implant size and insertion method could in theory have caused more damage to the neural tissue. Despite this, over the time points observed we found decreased FBR for the flexible electrodes compared to the microwires. This effect is most likely due to less irritation of the tissue by a mechanically compliant implant.

Although we do not report time points earlier than 26 weeks, electrophysiology obtained from the sinusoidal probe shows that the probe was reliable in obtaining high fidelity signals before this histology end point compared to the microwire. However, these timepoints for the FBR should also be evaluated in the future.

Single penetrations were made in this study and indeed during surgery we were able to angle electrodes slightly to miss prominent surface blood vessels, which would be detrimental to the signal quality obtained from the electrodes. This supports existing literature that fixed geometry arrays can unavoidably damage vasculature, causing varied and detrimental chronic recording performance between arrays [6,7,10], which in some cases have been compared to stroke-like lesions [7].

Inserting a microelectrode also can cause damage to the blood-brain-barrier (BBB). Damage to the BBB would lead to an unfavorable environment for neurons as homeostasis would be lost through ionic imbalances and through the promotion of neurodegenerative pathways, which would lead to neuronal death. As the sinusoidal probe was able to record stable neural activity and have an overall reduced FBR responses overtime, it might be indicative that damage to the BBB was reduced, although this needs to be confirmed in future studies through specific staining using IBA1 primary antibody [7,11]. The effects of the interaction of the peripheral and central immune response merits further investigation as recent findings have shown a direct link between the two, rendering the brain a non-immuno privileged organ [21].

Although we performed conventional histological methods for evaluating the FBR surrounding the probes, recently published novel methods should be used to evaluate the FBR surrounding the electrode. Such methods include tissue-clearing methods, where a larger part of the brain can be sectioned and observed as an intact structure [22–24]. Further, these methods may allow for more than one primary antibody to be used, allowing for more thorough investigation of the implant sites. Often, the glial scar consists of densely packed cells around the electrode implant. A method of great interest to ‘zoom’ in on these cells and observe the interactions of the glial mediators and neurons is that of expansion microscopy. Using conventional histology methods and microscopy, it may be possible to closely examine the FBR by simply expanding the tissue and observing super high-resolution histology using conventional microscopes [25]. The findings could lead to new insights informing overall device design and a better understanding of the FBR.

We report data from a small number of animals, however our statistical comparisons were made by inserting a large number of electrodes to reduce animal to animal variations. However, more small animals and non-human primates should be implanted to corroborate these initial findings for the suitability of a flexible probe to enhance recording longevity through potential reduced micromotion related trauma.

## 7. Conclusions

Here we performed a comparison between a novel flexible microelectrode and conventional microwires over 26-96 week indwelling periods in the rabbit cortex. The flexible electrode was associated with decreased FBR and increased neural density across majority of the probe depth. These results suggest that flexible microelectrode designs help to reduce micromotion-induced trauma and enhance electrode longevity.

